# The role of feedback for sensorimotor decisions under risk

**DOI:** 10.1101/2025.07.03.662995

**Authors:** Christian Wolf, Artem V. Belopolsky, Markus Lappe

## Abstract

For goal-directed movements like throwing darts or shooting a soccer penalty, the optimal location to aim depends on the endpoint variability of an individual. Currently, there is no consensus whether people can optimize their movement planning based on information about their motor variability. Here, we tested the role of different types of feedback for movement planning under risk. We measured saccades towards a bar that consisted of a reward and a penalty region. Participants either received error-based feedback about their endpoint or reinforcement feedback about the resulting reward. We additionally manipulated the feedback schedule to assess the role of feedback frequency and whether feedback focusses on individual trials or a group of trials. Participants with trial-by-trial reinforcement feedback performed best and had the least endpoint deviation from optimality. Our results are consistent with a slow gradual drift in the participants’ internal aiming location under summary feedback. Feedback focusing on individual trials reduces this drift, thereby enabling consistent movement planning from one trial to the next. Our results therefore suggest that reinforcement feedback about a single movement is most effective to optimize movement planning under risk.

## Introduction

Even if we intend to perform the same movement multiple times, the movement and its outcome will not be identical, but it will vary somewhat from one time to the next. Consider, for example, shooting a penalty in soccer. To succeed, the ball must move towards the goal and overcome the goalkeeper. The latter is more likely when the ball is aimed close to the goal post. But even if we always aim for the lower left corner, our shot will sometimes miss the goal or hit the goal post instead, whereas in other cases the ball will land at a more central location of the goal. In goal-directed aiming tasks like shooting a penalty or throwing a dart, the optimal location to aim for, is determined by the motor variability of an individual. An experienced soccer player may aim close to the goal post, because he/she is aware that the ball will land close to the targeted location most of the time. A more unexperienced player, however, whose shooting is more variable, may repeatedly fail when aiming for the same location, because the ball will frequently hit the goal post or even land outside of the goal – reducing the chance to score to zero. The unexperienced player therefore benefits from aiming at a less extreme location (i.e., closer to the goal center).

Sensorimotor decision tasks like shooting a penalty or throwing darts can be considered a case of movement planning under risk, and performance in these tasks depends on sensory uncertainty, motor uncertainty as well as the reward structure of the task (for reviews see Trommershäuser et al., 2008; Wolpert & Landy, 2012). The reward structure refers to the magnitude of reward and penalty as well as the relative size of the reward and penalty region. Consequently, the reward structure can be manipulated, for example, by changing the relative size of the reward and penalty region, or by imposing a different ratio between reward and penalty. Sensory and motor uncertainty on the other hand, can be considered a characteristic of the participant performing the task. Previous research has shown that movement planning under risk considers both, the reward structure of the task as well as the inherent motor variability (Trommershäuser et al., 2003a, 2003b; Zhang et al., 2013). Moreover, when feedback about endpoints was perturbed, thereby increasing the inferred motor variability, people adjust their behavior and select a less risk-seeking point to aim (Trommershäuser et al., 2005). Whereas movement planning has been shown to maximize the expected gain and was therefore considered close to optimal (Trommershäuser et al., 2003b, 2003a, 2005), other studies reported movement planning to be sub-optimal (Ota et al., 2016, 2019; Wu et al., 2006). For example, Wu et al. (2006) showed that behavior can be sub-optimal with more complex, asymmetric reward structures. But even for simpler reward structures, behavior can be sub-optimal, even after prolonged training (Ota et al., 2016) or when additional information was provided to the participant (Ota et al., 2019). To this end, Ota et al. (2019) provided participants with blocked summary feedback after a block of 50 trials. This feedback showed the endpoints of the 50 recent trials, reasoning that observing the distribution of many trials helps to directly infer motor variability. Yet, the additional summary feedback did not improve performance (Ota et al., 2019). However, given that summary feedback was provided incrementally, it is unclear whether blocked summary feedback can improve movement planning at all or whether it is as effective as trial-by-trial feedback.

In the literature on motor learning and skill acquisition, there is an ongoing debate about the effectiveness of feedback with a reduced frequency compared to trial-by-trial feedback (Fujii et al., 2016; Marco-Ahulló et al., 2024; McKay et al., 2022; Ronsse et al., 2011; Salmoni et al., 1984; Schmidt et al., 1989; Weir-Mayta et al., 2022; Wulf et al., 2010). According to the guidance hypothesis (Salmoni et al., 1984; Schmidt et al., 1989), trial-by-trial feedback can be superior during skill acquisition but can be detrimental once the feedback is removed, because participants are too reliant upon the presence of feedback. Ever since empirical studies have provided evidence for and against the guidance hypothesis. A recent meta-analysis (McKay et al., 2022) found no clear evidence for the guidance hypothesis and superior retention performance with reduced feedback frequencies. Moreover, trial-by-trial and summary feedback not only differ in terms of feedback frequency but also in terms of the focus of feedback: Trial-by-trial feedback provides information about a single movement or its outcome whereas summary feedback provides information about multiple movements. For movement planning under risk a focus on multiple movements appears beneficial, because it allows to directly assess the variability of the movement – and thus the crucial property to maximize performance. In contrast to that, neurocomputational accounts of human motor learning emphasize the evaluation of individual movements and their outcomes (Diedrichsen et al., 2010; Doya, 2000; Feulner et al., 2025; Krakauer & Mazzoni, 2011; Shadmehr et al., 2010), suggesting better performance when individual movements can be evaluated.

Motor learning is supported by at least two different mechanisms: error-based learning and reinforcement learning. Error-based learning relies on sensory prediction errors – the discrepancy between expected and actual sensory outcomes of a movement – and enables precise, direction-specific corrections based on this mismatch (Shadmehr et al., 2010). In contrast, reinforcement learning uses reward prediction errors to adjust future actions based on their success or failure, without requiring explicit knowledge about the movement error direction (Doya, 2000; Izawa & Shadmehr, 2011). These two learning systems differ in their computational principles but also in their neural substrates, with the cerebellum primarily involved in error-based learning (Catz et al., 2008; Herzfeld et al., 2018; Marr, 1969) and the basal ganglia and the dopaminergic system playing a central role in reinforcement learning (Hikosaka et al., 2014; Schultz, 1998). While both learning mechanisms have been extensively studied in motor adaptation, less is known how they contribute to sensorimotor decisions under risk and thus in situations in which not only movement accuracy, but also variability and the reward structure needs to be considered.

In the present study, we rigorously tested the role of feedback for movement planning under risk using saccade eye movements – a response system with established influences of reinforcement and error-based learning (Madelain et al., 2011; Pélisson et al., 2010). We asked participants to make saccades to an elongated bar that was divided in a reward and a penalty region. Both, reward and penalty, increased toward the center of the bar. Thereby, aiming into the rewarded region close to the center of the bar yields a high amount of reward – yet at the risk of encountering a penalty. We systematically manipulated the feedback modality (error-based feedback, reinforcement feedback) as well as the feedback schedule (trial-by-trial feedback, blocked summary feedback, rolling summary feedback) in a fully crossed between-participant design. Participants either received feedback about the obtained score or about their endpoints, enabling either reinforcement or error-based learning. Moreover, participants either received feedback on a trial-by-trial level or they received feedback about the recent 30 trials (summary feedback). Two groups of participants received summary feedback every 30 trials (blocked summary) whereas two other groups received summary feedback after every trial (rolling summary). A comparison between the blocked summary and rolling summary groups reveals the contribution of feedback frequency, and the comparison between trial-by-trial and rolling summary feedback reveals the contribution of feedback focus – thus, whether feedback focusses on the outcome of individual movements or on a set of movements.

## Methods

### Participants

We recorded data of 120 participants (100 females, 20 males; age range: 18–46; median age: 21). Participants were Psychology students from the University of Muenster and were reimbursed with course credit or with 8€/h. Additionally, participants received a performance-contingent reward that depended on their performance in the experiment. The performance-contingent reward varied between 0€ and 2.40€ (median: 1.90€). Participants had normal or corrected-to-normal vision. The experiment was approved by the Ethics Committee of the Faculty of Psychology and Sports Science at the University of Münster and participants provided written informed consent before taking part in the experiment.

### Setup and stimuli

Stimuli were presented on an Eizo FlexScan 22-inch CRT monitor (Eizo, Hakusan, Japan) with a resolution of 1152 × 870 pixels, an effective display size of 40.7 × 30.5 cm and a refresh rate of 75 Hz. A chin–forehead rest was used to restrict head movements and assure a viewing distance of 67 cm. Stimulus presentation was controlled via the Psychtoolbox (Brainard, 1997; Kleiner et al., 2007) in MATLAB (The MathWorks, Natick, MA). Eye position of the right eye was recorded at 1000 Hz using the EyeLink 1000 (SR Research, Mississauga, ON, Canada) and the EyeLink Toolbox (Cornelissen et al., 2002). All stimuli were presented on a gray background (9.06 cd/m^2^). The EyeLink was calibrated at the beginning and in the middle of each block using a 9-point calibration protocol.

The stimulus was a horizontally elongated white bar (83.6 cd/m^2^) with a width of 8.1 deg and a height of 1.04 deg. It was presented at the horizontal monitor midline, at a vertical eccentricity of either -7.5 deg or +7.5 deg. The bar had no sharp visible edges but slowly faded into the background: The outmost 0.15 deg of every edge were a linear transition into the background. We used a combination of fixation cross and bull’s eye as fixation cross (Thaler et al., 2012). To provide feedback in the error-based conditions, we used an ellipsoid with a diameter of 0.3 (horizontal) and 0.5 (vertical) for trial-by-trial feedback and black vertical lines of 1.65 deg length for summary feedback (Fig. 1A). For the reinforcement conditions, we used numbers rounded to one decimal displayed 1.25 deg above the bar.

**Figure 1.**
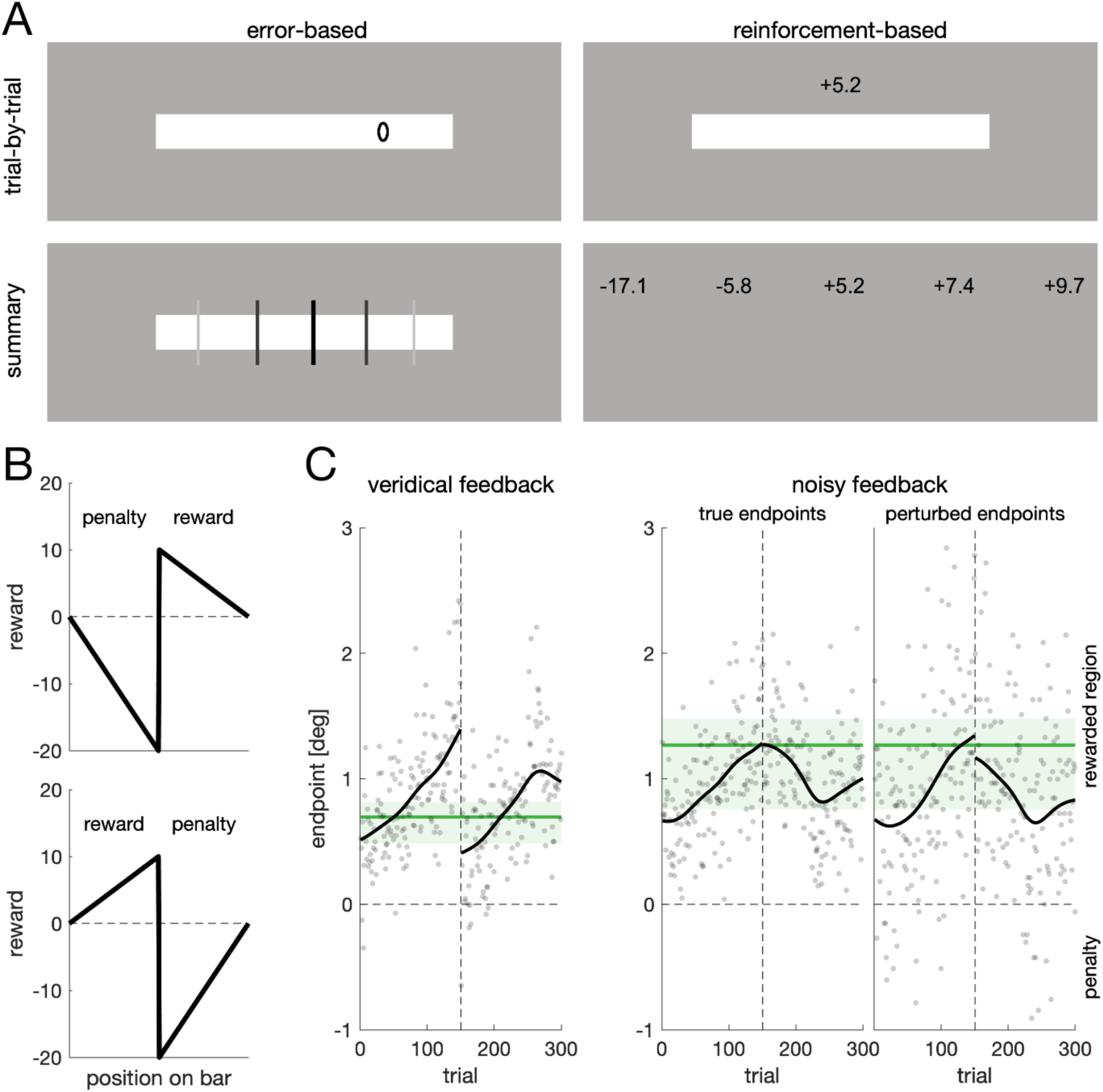
Design and critical manipulations. (**A**) Participants were rewarded/penalized for making vertical eye movements to a horizontally elongated white bar. We recorded six groups that received different forms of feedback. Participants either received feedback about their endpoint (error-based, left column) or about their obtained score (reinforcement, right column). Participants in the trial-by-trial groups (top row) received feedback about their performance in the recent trial, whereas participants in the summary groups (bottom row) received feedback about their performance in the recent 30 trials. Participants in the blocked summary groups received feedback every 30 trials, participants in the rolling summary group received feedback after every trial. Summary feedback consisted of the mean, the minimum and maximum, as well as the 25^th^ and 75^th^ percentile either in terms of position or obtained score. (**B**) Reward structure. One half of the bar was associated with a reward, the other half with a penalty. Importantly, reward and penalty increased towards the bar center. For one half of participants the rewarded region was the righthand part of the bar. (**C**) Individual data of one participant. Every participant completed two blocks, each with a break after 150 trials (dashed vertical line). In the first block participants received feedback about their true endpoints (veridical feedback, left panel). In block two (noisy feedback, center and right panel) position noise was added to the provided feedback. Every data point is the endpoint of one trial, solid black lines are a moving average (*σ* = 20 trials). Green lines are optimal aim points derived from the variability relative to the moving average. The green shaded area denotes the region of optimality for that individual (Suppl. Fig. S1). For noisy feedback, the perturbed endpoints (right panel) and not the true endpoints (center panel) were used to compute the optimal aim point.

### Design and Procedure

Participants were instructed to make vertical saccades to a horizontally elongated bar. Half of the bar was associated with a reward, whereas the other half was associated with a penalty. Critically, reward and penalty increased towards the center of the bar, with the penalty being twice as high as the reward (Fig. 1B). This information was made explicit to participants. The vertical position of the bar and the reward orientation on the bar were balanced across participants. The asymmetric reward structure ensured that the expectedly few penalties contributed to the overall performance. We recorded six groups that differed in terms of the feedback provided to them. Participants either received feedback about their endpoint on the bar (error-based), or about the obtained reward (reinforcement). Moreover, participants received either feedback about the recent trial (trial-by-trial) or about the recent 30 trials (blocked summary and rolling summary). Participants in the blocked summary groups received feedback every 30 trials, whereas participants in the rolling summary groups received feedback after every trial. During the first 29 trials, participants in the rolling summary conditions received feedback about all previous trials.

Every participant completed two blocks. In the first block participants received feedback about their true endpoints (veridical feedback). In the second block (noisy feedback) endpoints (and thus scores) were perturbed by adding position noise (*σ* = 0.5 deg) to the true endpoint. Thus, the experiment constituted a 2 × 2 × 3 design, with the within-participant factor feedback veridicality (veridical, noisy) and the between-participant factors feedback modality (error-based, reinforcement) and feedback schedule (trial-by-trial, blocked summary, rolling summary).

At the beginning of each trial a fixation cross was displayed at the screen center. A trial was started if gaze was within a square region of 2.5 deg width around the fixation cross for 250 consecutive samples. After a uniform random interval between 200 and 500 ms, the bar appeared. Participants were instructed to quickly look at the bar after its appearance by means of a single eye movement. A sample was labelled on-target if it was less than 0.5 deg away from the edge of the bar. The target was classified as selected if gaze was on the bar for 200 samples. The mean gaze position of the last ten samples was used to provide feedback. In the trial-by-trial conditions, feedback was displayed for 306 ms. Summary feedback was displayed until participants pressed any button on a keyboard. If the target was not selected within 1200 ms, the target was removed, and the next trial began. Summary feedback was a graphical representation (Fig. 1A) of the mean, the minimum and maximum, as well as the 25^th^ and 75^th^ percentile either in terms of position or score. Participants in the summary groups were told that feedback provided information about the recent 30 trials, displaying the mean (central line or value; see Fig. 1A), the values covering the central half of the responses (neighboring elements) as well as the minimum and maximum (outmost elements).

Every participant completed a demo version of the experiment before proceeding to the main experiment to get familiar with the task. The demo consisted of five trials and no feedback was provided. Unlike the main experiment, the penalized region of the bar was displayed in dark gray. Both blocks of the main experiment consisted of 300 trials and included a break after 150 trials. Participants were told that their task was the same in the two blocks, yet that a different algorithm was used to compute where they had looked. After completing the second block, score points were converted into a monetary reward, with 1000 points yielding 1€. For the noisy feedback block, the payment was either based on the true score or on the perturbed score – depending on what was higher.

### Data analysis

We recorded gaze position of the right eye at 1000Hz. Saccade onsets and offsets were defined using the EyeLink 1000 algorithm. In the offline analysis, gaze position at saccade offset was used as saccade endpoint. These offline estimates of saccade endpoints and the online estimates of gaze position (which were used for feedback) were highly correlated, r = 0.96, p < 0.001. The analysis is based on the offline results. We discarded 21 trials (out of 72000) because no gaze position on the target was detected.

The optimal aiming point depends on the motor variability. Under the assumption of a stable aim point across trials, the motor variability is identical to the variability of endpoints. An optimal aim point can then be derived by shifting the entire endpoint distribution to different mean values and computing the resulting overall score for that mean endpoint. The mean endpoint with the highest score can be used as an estimate of the optimal aiming point. However, such an analysis would neglect that participants may change their aiming location over the course of the experiment, either by slow gradual changes or by sudden changes in strategy. Thus, the observed variability may not only reflect motor variability but also aiming variability (i.e., variability due to aiming at different locations). In consequence, this approach would overestimate the individual motor variability. Alternatively, one may compute the endpoint residuals relative to a moving average (Fig. 1C). This way, changes in the aiming location over time can be accounted for. However, the moving average and the resulting residuals will depend on the size of the sliding window that is used to compute the moving average. A small sliding window may result in a noisy moving average that underestimates motor variability. A large sliding window, on the other hand, will overestimate motor variability, given that the moving average will approach the individual mean with increasing window size. Considering all these limitations, we decided to not provide point estimates of motor variability and thus of optimal performance, but to compute an upper limit (spread around the individual mean) and a lower limit of motor variability (spread around the moving average with the smallest sliding window). We refer to the region in between the upper and the lower limit as region of optimality (Suppl. Fig. S1). Specifically, to compute the lower limit of motor variability, we used a sliding Gaussian window with a standard deviation of 1 trial. For Figures 2–4, the depicted regions of optimality indicate the distance from the lower limit (minus 1.96 times the standard error) to the upper limit (plus 1.96 times the standard error) for the respective groups of participants. We excluded one participant from the computation of the region of optimality, because this participant alternated between looking at the left and right edge of the bar during the first block, thereby producing unrealistic high values of variability.

**Figure 2.**
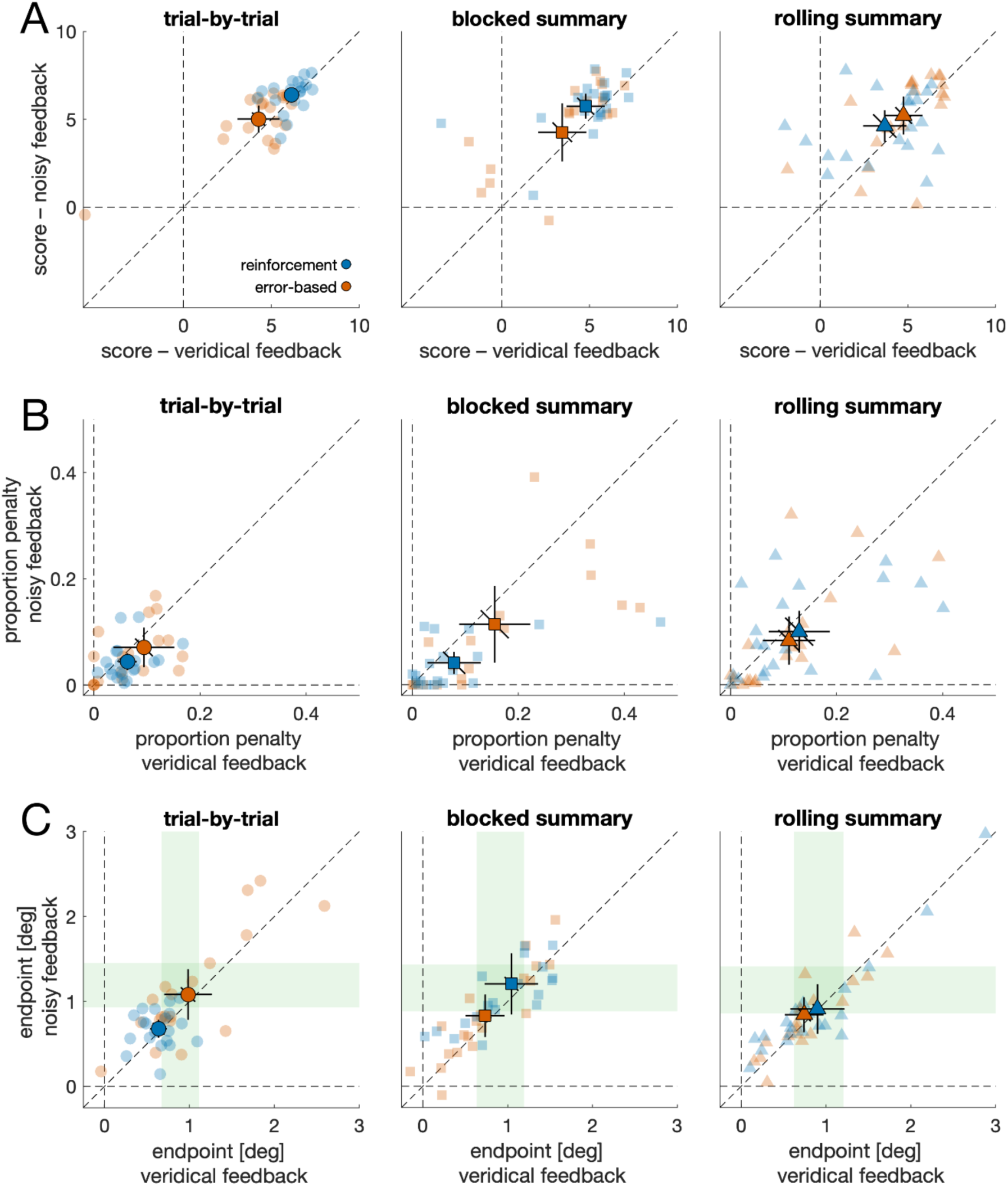
Performance. (**A**) Average score per trial for every participant. Every participant first completed a veridical feedback condition before proceeding to the noisy feedback condition. The score in the noisy feedback condition shows the true score – not the perturbed one used for feedback. Each faint data point is the mean of one individual. Solid data points are the group mean. Error bars represent the 95% confidence interval of between-participant variability. Diagonal error bars must be compared to the identity line. (**B**) Proportion penalized trials for every participant. (**C**) Average endpoint of every participant. Positive values denote average endpoints on the rewarded region; negative values indicate the penalized region. Green regions denote the region of optimality.

We compared scores and endpoints using a 2 × 2 × 3 ANOVA with the within-participant factor feedback veridicality (veridical, noisy) and the two between-participant factors feedback modality (error-based, reinforcement) and feedback schedule (trial-by-trial, blocked summary, rolling summary). The direction of effects was tested using t-tests. Inferential statistics are supplemented with effect size estimates, Cohen’s d for t-tests, and partial eta squared, *η*²_*p*_, for ANOVA results.

We used cluster-permutation tests with 1000 permutations to compare continuous data (i.e., endpoints over time and kernel-density estimates) of two conditions. For each permutation, the assignment of participants to the groups was randomly permuted.

## Results

### Best performance with trial-by-trial reinforcement feedback

To quantify performance, we computed the mean score per trial. Figure 2A shows scatter plots of the noisy score (i.e., score in the noisy feedback condition) over the veridical score (i.e., score in the veridical condition) for the different groups. To compute the noisy score, we used the true score associated with the real endpoints and not the perturbed score that was used for feedback.

Scores were compared with a 2 × 2 × 3 ANOVA. On average, scores were higher with reinforcement feedback compared to error-based feedback, *F*(1,114) = 4.01, *p* = 0.048, *η*²_*p*_ = 0.034. Crucially, the effect of feedback modality was further modulated by feedback schedule, *F*(2,114) = 4.63, *p* = 0.012, *η*²_*p*_ = 0.075: Whereas reinforcement feedback was superior to error-based feedback when provided on a trial-by-trial basis, veridical: *t*(38) = 3.08, *p* = 0.004, *d* = 0.97; noisy: *t*(38) = 3.22, *p* = 0.003, *d* = 1.02, no such difference could be observed with blocked summary feedback, veridical: *t*(38) = 1.60, *p* = 0.119, *d* = 0.50; noisy: *t*(38) = 1.73, *p* = 0.092, *d* = 0.55, or with rolling summary feedback, veridical: *t*(38) = -1.39, *p* = 0.174, *d* = -0.44; noisy: *t*(38) = -0.89, *p* = 0.382, *d* = -0.28. Moreover, for reinforcement feedback, scores were higher for trial-by-trial feedback compared to rolling summary feedback, veridical: *t*(38) = 3.97, *p* < 0.001, *d* = 1.26; noisy: *t*(38) = 3.70, *p* < 0.001, *d* = 1.17, as well as compared to blocked summary feedback, but only in the veridical condition, veridical: *t*(38) = 2.43, *p* = 0.0198, *d* = 0.77; noisy: *t*(38) = 1.58, *p* = 0.122, *d* = 0.50. Additionally, the ANOVA revealed a main effect of feedback veridicality, reflecting that noisy scores were higher than veridical scores, *F*(1,114) = 13.694, *p* < 0.001, *η*²_*p*_ = 0.107. This is expected if, first, our manipulation of feedback veridicality was successful (i.e., people becoming more cautious with noisy feedback) and, second, participants lose most points in the veridical condition by being too risky.

Performance in the task depends on aiming at a rewarded location close to the center of the bar while minimizing the number of penalties. A low score can result from being too cautious (low reward, few if any penalties) or from being too risky (high reward, but also high number of penalties). Indeed, over all groups, the mean score was strongly correlated with the proportion of trials in the penalty region, *r*(238) = -0.85, *p* < 0.001. Descriptively, the proportion of penalties was lowest with reinforcement feedback, both in the trial-by-trial and blocked summary groups (Fig. 2B). Yet, the ANOVA only yielded a main effect of feedback veridicality, *F*(1,114) = 14.89, *p* < 0.001, *η*²_*p*_ = 0.116, reflecting a lower number of penalties in the noisy condition (unperturbed data).

### Trial-by-trial reinforcement feedback enables high reward endpoints

The relationship between the individual mean endpoint on the bar and the mean score per trial was inverted U-shaped (Fig. 3) with a peak at approximately 1 deg (veridical feedback condition): For endpoints above 1 deg, the score was lower the further the mean endpoint was away from the bar center, *r*(34) = -0.75, *p* < 0.001, whereas the opposite was true for endpoints below 1 deg, *r*(82) = 0.67, *p* < 0.001. This turning point of approximately 1 deg coincides with the inferred region of optimality: In the veridical condition, the lower limit of optimality was *M_lower_* = 0.69 deg (*SD_lower_* = 0.14 deg), and the mean upper limit was *M_upper_* = 1.10 deg (*SD_upper_* = 0.24 deg). In the noisy condition, however, the lower limit was *M_lower_* = 0.93 deg (*SD_lower_* = 0.12 deg) and the upper limit was *M_upper_* = 1.38 deg (*SD_upper_* = 0.18 deg). Thus, across the whole sample, we obtained a region of optimality that varied between 0.66 deg and 1.14 deg for the veridical and between 0.90 deg and 1.41 deg for the noisy condition. Please note that the regions of optimality are not identical for the different groups (Figs. 2C & 4A).

**Figure 3.**
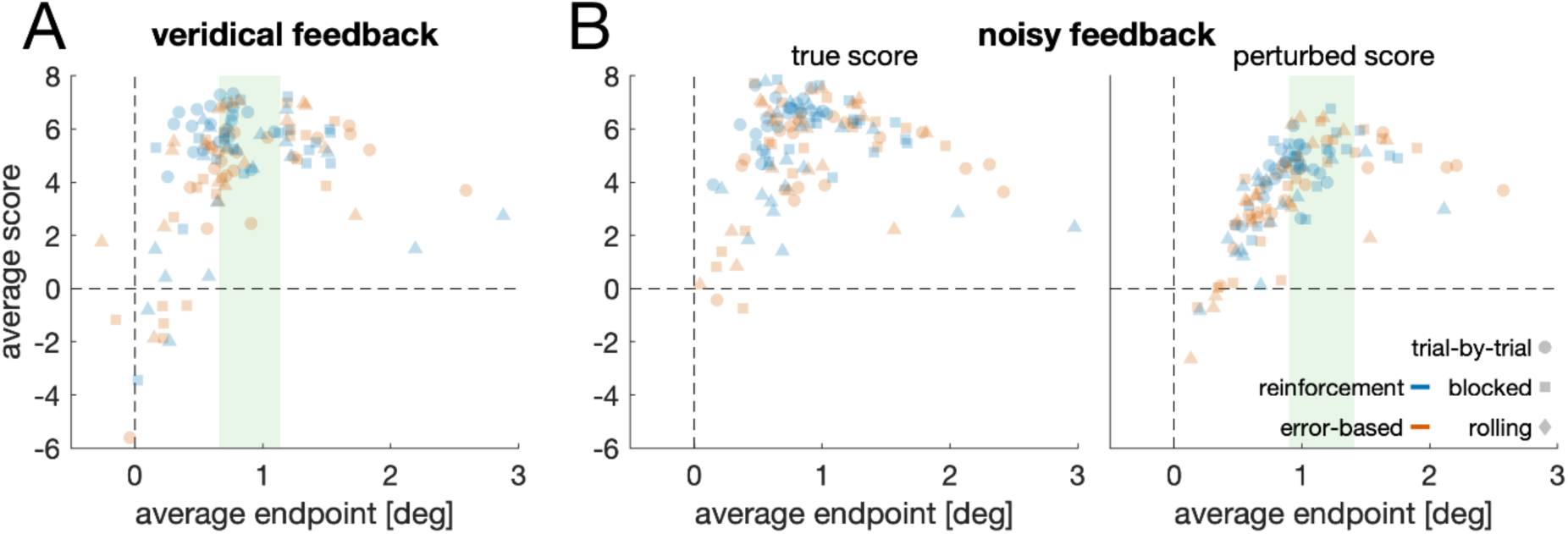
Scores as a function of endpoints. Average score as a function of the average endpoint for (**A**) the veridical feedback condition and (**B**) the noisy feedback condition. Each data point is one individual. Green regions denote the regions of optimality.

**Figure 4.**
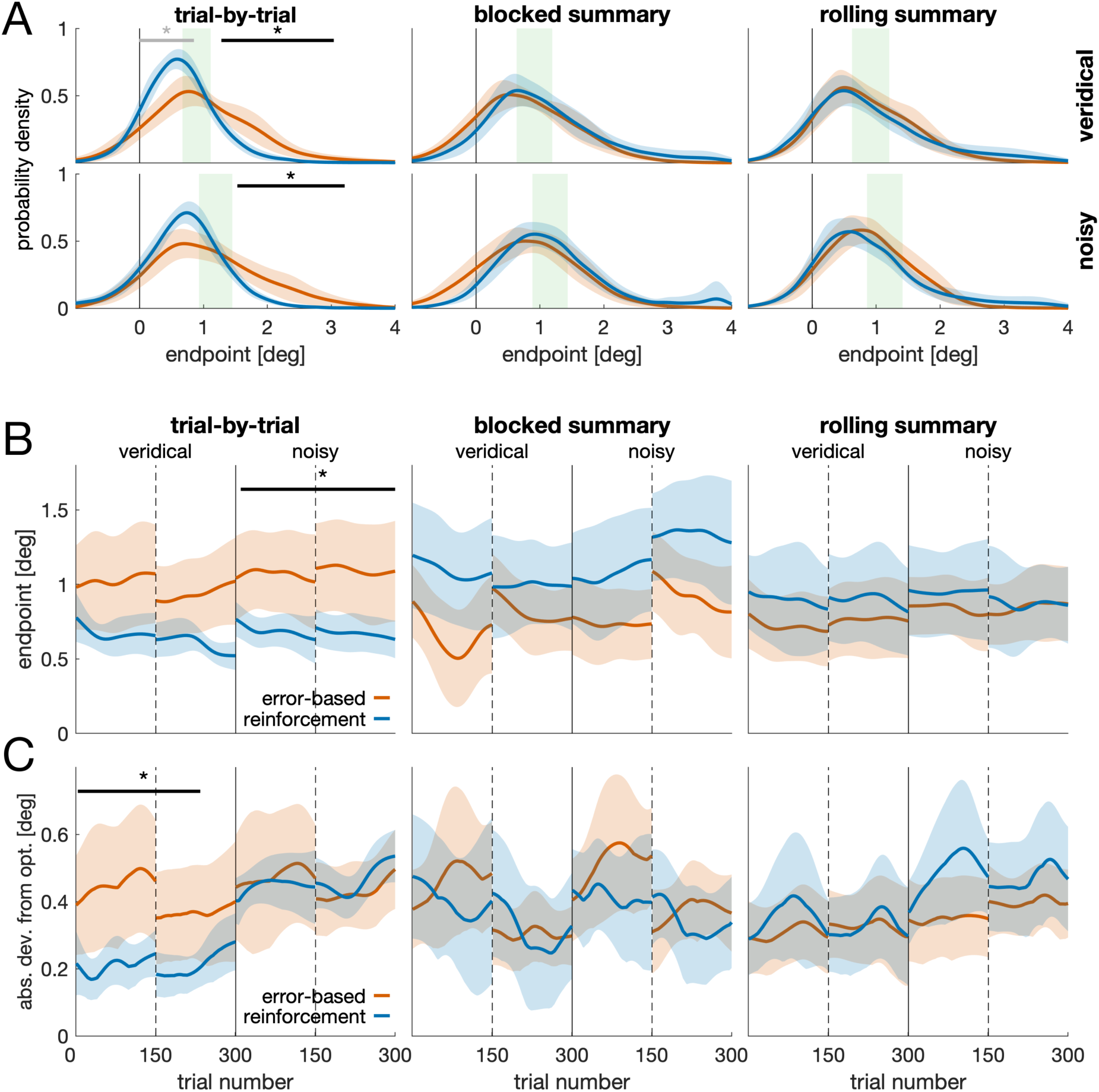
Endpoints in space and time. (**A**) Kernel density estimates of saccade endpoints. Horizontal lines and asterisks denote a difference between error-based feedback (orange) and reinforcement feedback (blue) as revealed by a cluster-permutation test. Black colors indicate the strongest cluster. (**B**) Endpoints over time for error-based (orange) and reinforcement feedback (blue), separate for the trial-by-trial (left panel), blocked summary (center panel) and rolling summary condition (right panel). Each panel shows data from the veridical feedback condition (left of solid vertical line) and the noisy feedback condition (right of solid vertical line). Dashed vertical lines indicate the break during each experimental block. Solid horizontal lines and asterisks indicate a significant difference between time courses as revealed by a cluster-permutation test. (**C**) Absolute deviation from the region of optimality. Same conventions as in (B).

Whereas mean endpoints showed substantial interindividual differences in most of the groups (Fig. 2C), participants in the trial-by-trial reinforcement feedback group were most consistent (lowest between-participant variability; Fig. 2C) and their mean endpoints were either within the region of optimality or risk-seeking (i.e., in-between the region of optimality and the bar center; Fig. 4A). An ANOVA on the endpoints revealed an interaction between feedback modality and feedback schedule, *F*(1,114) = 4.70, *p* = 0.011, *η*²_*p*_ = 0.076. For trial-by-trial feedback, endpoints were lower (i.e., closer to the bar center) for the reinforcement versus the error-based condition, veridical: *t*(38) = 2.44, *p* = 0.020, *d* = 0.77; noisy: *t*(38) = 2.67, *p* = 0.011, *d* = 0.84. This was neither the case for blocked summary feedback, veridical: *t*(38) = - 1.67, *p* = 0.103, *d* = -0.53; noisy: *t*(38) = -1.79, *p* = 0.008, *d* = -0.57, nor for the rolling summary, veridical: *t*(38) = -0.84, *p* = 0.406, *d* = -0.27; noisy: *t*(38) = -0.38, *p* = 0.701, *d* = -0.12. For reinforcement feedback, endpoints were closer to the bar center for trial-by-trial compared to blocked summary feedback, veridical: *t*(38) = 2.59, *p* = 0.013, *d* = 0.82; noisy: *t*(38) = 2.94, *p* = 0.006, *d* = 0.93, but not compared to rolling summary, veridical: *t*(38) = 1.64, *p* = 0.109, *d* = 0.52; noisy: *t*(38) = 1.56, *p* = 0.127, *d* = 0.49.

Moreover, the ANOVA revealed a main effect of feedback veridicality, *F*(1,114) = 9.24, *p* = 0.003, *η*²_*p*_= 0.075, reflecting that endpoints in the noisy feedback condition, *M_noisy_* = 0.93 deg, *SD_noisy_* = 0.58 deg, were on average further away from the bar center than in the veridical feedback condition, *M_veridical_* = 0.84 deg, *SD_veridical_* = 0.56 deg, *t*(119) = 3.07, *p* = 0.003, *d* = 0.28. This reflects that the noisy feedback manipulation was successful and that participants adjusted their endpoints to become more cautious. Yet, this average shift (*ΔM* = 0.09 deg) was less than would be required to maintain a good level of performance, given that the differences in the lower limit (*ΔM* = 0.24) and upper limit of the optimal endpoint (*ΔM* = 0.28) were approximately three times as high.

In a next step, we analyzed the spatial (Fig. 4A) and temporal distribution (Fig. 4B-C) of endpoints. Based on the individual region of optimality, we computed the proportion of rewarded but sub-optimal trials and classified them as either risk-seeking (i.e., in between bar center and region of optimality) or as cautious/loss-aversive (i.e., beyond the region of optimality). For five out of six groups the fraction of risk-seeking and loss-aversive trials was approximately the same (Fig. 4A; Suppl. Table S1), with trial-by-trial reinforcement feedback being the only exception. With veridical feedback, participants in the trial-by-trial reinforcement group had a higher proportion of risk-seeking trials, *M_risk_* = 46.7% [42.8% 50.6%] and a lower number of cautious/loss-aversive trials, *M_cautious_* = 22.7% [17.0% 28.4%], compared to the remaining sample, *M_risk_* = 30.1% [26.9% 33.2%] and *M_cautious_* = 35.5% [30.3% 40.8%] respectively. This was also true for noisy feedback: *M_risk_* = 58.9% [53.8% 64.1%] and *M_cautious_* = 12.9% [9.0% 16.8%] for trial-by-trial reinforcement feedback compared to *M_risk_* = 41.1% [36.7% 45.5%] and *M_cautious_* = 27.8% [22.6% 33.3%] for the remaining sample. For each individual, we estimated the endpoint distribution using a kernel smoothing function and compared distributions of different groups using cluster-permutation tests (Fig. 4A). For trial-by-trial feedback, we observed a difference between reinforcement and error-based feedback, veridical: *t_sum_* = 475.84, *t_crit_* = 200.35, *p* = 0.001, cluster position: 1.28 – 3.04 deg; noisy: *t_sum_* = 428.54, *t_crit_* = 232.95, *p* = 0.008, cluster position: 1.53 – 3.21 deg.

How much do endpoints deviate from optimality? Based on the time course of endpoints (Fig. 4B), we computed the distance of each endpoint to the region of optimality of an individual. Figure 4C shows a moving average of this deviation. The deviation to optimality was lowest with trial-by-trial reinforcement feedback in the veridical condition. A cluster-permutation test revealed a difference between reinforcement and error-based feedback, *t_sum_* = 560.9, *t_crit_* = 261.9, *p* = 0.003, trials: 4 – 234.

### Trial-by-trial feedback enables immediate error correction

How does movement planning change after making an error? Figure 5A shows endpoints (relative to the individual mean) of the five trials (T1 to T5) following a saccade into the penalty region. Whereas endpoints immediately following a penalty are not different from the individual mean with trial-trial-feedback, *M_T1_* = -0.04 deg, *t*(36) = 0.98, *p* = 0.332, *d* = 0.16, endpoints immediately following an error remain below the individual mean, both for blocked summary feedback, *M_T1_* = -0.34, *t*(38) = 6.90, *p* < 0.001, *d* = 1.10, and for rolling summary feedback, *M_T1_* = -0.17, *t*(38) = 3.54, *p* = 0.001, *d* = 0.57. The endpoints in both summary conditions do appear to show a gradual return to the individual mean (Fig. 5A). We compared post-penalty endpoints with a 2 × 3 × 5 ANOVA with the between-participant factors feedback modality and feedback schedule and the within-participant factor post-penalty trial (T1 to T5). Post-penalty behavior differed depending on the feedback schedule, *F*(2,109) = 14.05, *p* < 0.001, *η*²_*p*_ = 0.205, reflecting larger deviations from the individual mean for blocked summary feedback. Crucially, however, we found an interaction between post-penalty trial number and feedback schedule, *F*(8,436) = 2.66, *p* = 0.007, *η*²_*p*_ = 0.046, suggesting that the post-penalty time courses (Fig. 5A) are different for the different feedback schedules. Linear regressions fitted to the post-penalty trials of individuals showed a positive slope for blocked summary, *M_slope_* = 0.034, *t*(38) = 4.22, *p* < 0.001, *d* = 0.95, and for rolling summary, *M_slope_* = 0.029 *t*(38) = 2.13, *p* = 0.040, *d* = 0.48, but not for trial-by-trial feedback, *M_slope_* = -0.012, *t*(36) = -1.23, *p* = 0.227, *d* = -0.29. This reflects that endpoints immediately return to the individual mean with trial-by-trial feedback, whereas they show a more gradual return with summary feedback.

**Figure 5.**
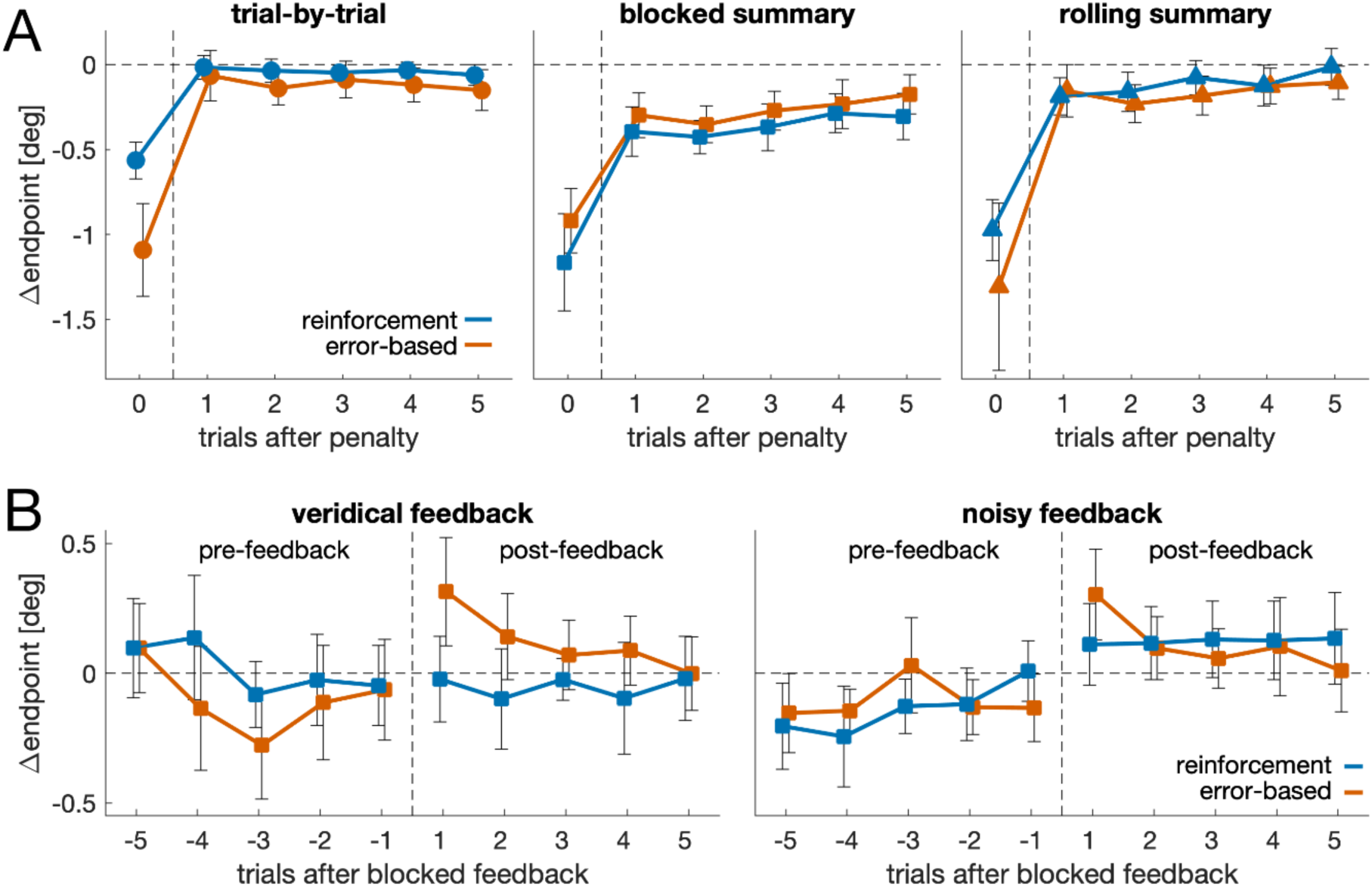
Post-error and post-feedback behavior. (**A**) Endpoints following a saccade on the penalized region for trial-by-trial feedback (left panel), blocked summary feedback (central panel) and rolling summary feedback (right panel). Endpoints are shown relative to the individual mean (horizontal dashed line). The results in (A) and the accompanying analysis combines data from the veridical and noisy feedback condition. The pattern and conclusions do not change if the analysis was restricted to data from the veridical block. (**B**) Endpoints before and after feedback in the blocked summary condition. Endpoints are shown relative to the individual mean. All error bars are 95% confidence intervals of between-participant variability.

An immediate return to the individual mean (from T0 to T1) with a subsequent (T1 to T5) slope of (approximately) zero, as it is found with trial-by-trial feedback, does not necessarily reflect an active error-correction process, but may instead reflect a regression to the mean. Under summary feedback, the return to the individual mean after an error is incomplete and unfolds gradually over time. Is the gradual return the default pattern that can also be observed in the absence of any feedback? To test this, we repeated the same analysis for the blocked summary condition, this time only selecting trials during which no feedback was displayed between T0 and T5 (Suppl. Fig. S2). Also in this selection, endpoints immediately following an error (T1), *M_T1_* = -0.34, were different from error trials (T0), *t*(38) = 7.86, *p* < 0.001, *d* = 1.49, as well as different from the individual mean, *t*(38) = 6.7, *p* < 0.001, *d* = 1.52. Most importantly, endpoints again showed a gradual return to the individual mean, *M_slope_* = 0.029, *t*(38) = 3.53, *p* = 0.001, *d* = 0.80. This shows that the gradual return that can be observed in the summary feedback groups (Fig. 5A) does occur independent of feedback.

### Endpoint adjustment after blocked feedback

Is blocked summary feedback thus any effective in guiding movement planning? To test whether information provided by blocked summary feedback was used, we compared pre-feedback and post-feedback trials in the blocked summary condition (Fig. 5B). Post-feedback endpoints were higher (i.e. more to the right) than those in pre-feedback trials, *F*(1,38) = 9.58, *p* = 0.004, *η*²_*p*_ = 0.201. This suggests that participants became more cautious after blocked summary feedback. When directly comparing trials before and after blocked feedback, this was especially evident after error-based feedback in the noisy block, veridical: *t*(19) = 2.09, *p* = 0.050, *d* = 0.47, noisy: *t*(19) = 3.74, *p* = 0.001, *d* = 0.83, but not with reinforcement feedback, veridical: *t*(19) = 0.17, *p* = 0.87, *d* = 0.04, noisy: *t*(19) = 0.91, *p* = 0.376, *d* = 0.20.

## Discussion

We tested the role of feedback for movement planning under risk. We asked participants to make saccades to an elongated white bar that consisted of a reward and a penalty region while manipulating the feedback modality (reinforcement versus error-based) and the feedback schedule (trial-by-trial, blocked summary, rolling summary). The different schedules either differed in terms of feedback frequency (blocked summary versus rolling summary) or in terms of feedback focus (trial-by-trial versus rolling summary). Results showed a large variability between participants in terms of performance, spanning the whole spectrum of different strategies (risk-seeking, optimal, loss-aversive; Figs 2–4). Trial-by-trial reinforcement feedback enabled a high level of performance for all participants. This was achieved by selecting an aim point (i.e., mean endpoint) that was either close to optimal or risk-seeking.

Our results therefore provide evidence for the superiority of trial-by-trial feedback when performing sensorimotor decisions under risk. Most importantly, our results suggest that feedback is most effective when provided on a trial-by-trial basis and when it simultaneously focusses on the outcome of a single trial compared to focusing on summary statistics of a group of trials. This latter aspect of feedback focus may appear counterintuitive given that the decision where to aim depends on the variability of your own movement – which can be more easily estimated from summary feedback. We had decided to provide summary feedback (mean, range and quartiles) rather than displaying endpoints (error-based) or score points (reinforcement) of the recent 30 trials, because we reasoned that the latter would not allow a fair comparison between error-based and reinforcement feedback: Whereas multiple endpoints and their variability may be assessable at a glance in a scatter plot (error-based feedback), a display of 30 score points (reinforcement) may be overwhelming and may require a prolonged and more detailed processing to obtain the same information. However, as has been shown by Ota et al. (2019), displaying a distribution of endpoints does not improve performance beyond that of trial-by-trial feedback. Moreover, we knew that our sample, which consisted exclusively of Psychology undergraduates, was well familiar with means, quartiles and ranges (although these words were not used for the instructions, see Methods). Thus, we cannot rule out the possibility that summary feedback is even less effective in a sample that is less familiar with statistics and distributions.

According to the guidance hypothesis (Salmoni et al., 1984; Schmidt et al., 1989), feedback with a reduced frequency can be more effective in motor learning and skill acquisition. However, the hypothesis mainly addresses performance in a retention test without any feedback – and not performance during the initial skill acquisition. Given that we did not include a retention test (i.e., a phase without any feedback), we cannot test this core prediction of the guidance hypothesis. However, we consider it unlikely that the guidance hypothesis holds true for movement planning under risk. To perform well in the task at hand, one must be able to select an aiming location based on the own motor uncertainty. Once participants have learnt which location yields a high amount of reward (as it is the case with trial-by-trial feedback), performance is unlikely to deteriorate strongly. On the other hand, if participants have not learnt to select a profitable aiming location during an initial feedback phase (as it is the case for some participants of the summary groups), it is unlikely that they learn to do so in a retention task without any further feedback. Hence, we would expect that the superiority of trial-by-trial reinforcement feedback for movement planning under risk also manifests in an immediate or delayed retention test.

Our findings show that trial-by-trial feedback is more successful when it provides information about the outcome of the behavioral goal (reinforcement) compared to information about the outcome of the movement (error-based). Whereas a success or failure is immediately apparent with reinforcement feedback, translating feedback about one endpoint into success or failure requires an additional step. For example, for an endpoint close to the bar center, one needs to assess whether the displayed feedback was on the lefthand (penalty) or the righthand half (reward). In that case, participants would have to perform a line bisection task – that is governed by visual uncertainty. In our task visual uncertainty could have been reduced by providing a visual reference, either by highlighting the center of the bar or by having differently colored reward and penalty regions. Hence, we cannot rule out the possibility that participants in the error-based condition would have performed better when an additional visual reference had been provided. However, we believe it unlikely that the presence or absence of a visual reference can account for all differences between the error-based and the reinforcement group, considering that approximately one fourth of the trial-by-trial error-based group pursued a loss-aversive strategy.

Our results also show that people change their movement planning based on perturbed feedback to become more cautious. Yet, this adjustment was comparatively small and only covered approximately 25% of the adjustment that would have expected given the manipulation. One reason might be that participants attributed the larger errors in the second block externally – especially since this was emphasized in the instruction. Whereas oculomotor behavior can be adjusted to feedback indicating external errors (Heins & Lappe, 2024), this adjustment might have been incomplete (Gastrock et al., 2020; Wilke et al., 2013). Alternatively, this incomplete adjustment might reflect the sub-optimality in movement planning under risk (Ota et al., 2016; Wu et al., 2006). Here, we computed a region of optimality by computing an upper and a lower limit of optimality as opposed to computing point estimates. Hence, behavior outside that region can be clearly labelled sub-optimal, either as too risk-seeking or too cautious / loss-aversive. In almost all groups we observed both kinds of sub-optimality (Fig. 2C). One notable exception was the group receiving trial-by-trial reinforcement feedback, in which the deviation from optimal was lowest (Fig. 4C) and in which participants could either be classified as optimal or as risk-seeking. Thus, even in the condition resulting in the best performance we see an overall tendency for sub-optimality. One reason might be that people cannot represent asymmetric reward structures (Wu et al., 2006). In our task, the maximum penalty was twice as high as the maximum reward. Indeed, if the maximum penalty and reward had the same magnitude, the region of optimality would have been closer to the bar center (0.59 – 1.01 deg for the trial-by-trial reinforcement group in the veridical condition), resulting in larger overlap between behavior and optimality. Alternatively, sub-optimality might result from a need for certainty and occasional penalties reassure participants that they are aiming at a location yielding a high reward rate. The latter would predict that participants can perform well under an asymmetric reward, yet that they would rather do so when the maximum penalty is below the maximum reward – and not when it is the other way round as in the current task.

With trial-by-trial feedback, endpoints immediately returned to the individual mean after a penalty (Fig. 5A). This is consistent with a simple statistical assumption: if trial outcomes are sampled from a normal distribution around a fixed aim point, extreme values (e.g., penalties) will naturally be followed by less extreme values due to regression to the mean. In contrast, with summary feedback, the return to the mean is incomplete and unfolds gradually over several trials. One possible explanation is a slow drift in the participants’ internal aim point, driven by uncertainty about where exactly to aim. Without immediate feedback about an individual movement, participants may lose track of their intended target location, leading to a prolonged deviation across trials. Such drift would not only increase the overall endpoint variability (as observed in Fig. 4A) but also delay the recovery from a penalty region. This phenomenon may be particularly pronounced in tasks with spatially extended target and no additional visual reference. Thus, the immediate return to the individual mean with trial-by-trial feedback suggests that this type of feedback stabilizes the internal representation of the aim point and supports consistent movement planning from trial to trial.

To summarize, we here show that trial-by-trial reinforcement feedback is superior when performing sensorimotor decisions under risk. Participants with trial-by-trial reinforcement feedback were most consistent and selected an aim point yielding a high rate of reward. Crucially, trial-by-trial feedback reinforcement is effective because it conveys information about a single movement (feedback focus) and not because it is administered after every trial (feedback frequency). Our data are consistent with a slow drift in the participants’ internal aim point under summary feedback. Trial-by-trial feedback reduces this drift and enables consistent movement planning from one trial to the next.

## Acknowledgments

This work was funded by the Deutsche Forschungsgemeinschaft (DFG, German Research Foundation), Project No. 427754309, awarded to CW. The authors thank Linda Melsheimer, Max Schuhriemen, Josefina Dreiling and Linda Tackenberg for collecting the data. The authors have no competing interests to declare that are relevant to the content of this article.

