## Supplementary material for "The role of feedback for sensorimotor decisions under risk": Suppl. Table S1

**Supplementary Table S1. Endpoint classification**. Based on the individual region of optimality, endpoints were classified as penalty, risk-seeking (i.e., in between bar center and region of optimality), optimal or loss-aversive (i.e., beyond the region of optimality). Each cell shows the mean percentage of trials as well as the 95% confidence interval for a given condition.

| condition | | | percentage of trials | | | |
|---|---|---|---|---|---|---|
| feedback type | feedback schedule | feedback veridicality | penalty | risk-seeking | optimal | loss-aversive |
| rein-forcement | trial-by-trial | veridical | M = 10.4% [7.2% 13.6%] | M = 46.7% [42.8% 50.6%] | M = 20.2% [16.9% 23.5%] | M = 22.7% [17.0% 28.4%] |
| | | noisy | M = 12.3% [7.5% 17.0%] | M = 58.9% [53.8% 64.1%] | M = 15.8% [11.1% 20.4%] | M = 12.9% [9.0% 16.8%] |
| | blocked summary | veridical | M = 8.7% [3.2% 14.1%] | M = 26.2% [20.3% 32.1%] | M = 22.3% [17.6% 26.9%] | M = 42.8% [31.4% 54.1%] |
| | | noisy | M = 4.7% [2.5% 6.9%] | M = 37.5% [28.5% 46.5%] | M = 22.2% [16.7% 27.8%] | M = 35.6% [23.7% 47.5%] |
| | rolling summary | veridical | M = 13.6% [8.1% 19.1%] | M = 33.0% [26.9% 39.1%] | M = 23.0% [18.3% 27.7%] | M = 30.3% [20.0% 40.5%] |
| | | noisy | M = 10.9% [7.6% 14.1%] | M = 47.7% [38.6% 56.7%] | M = 18.9% [13.9% 23.8%] | M = 22.6% [12.1% 33.2%] |
| error-based | trial-by-trial | veridical | M = 10.5% [4.4% 16.6%] | M = 28.9% [20.9% 37.0%] | M = 20.8% [15.0% 26.6%] | M = 39.7% [26.4% 53.0%] |
| | | noisy | M = 8.9% [4.3% 13.5%] | M = 39.0% [28.3% 49.7%] | M = 18.7% [14.1% 23.3%] | M = 33.4% [19.4% 47.4%] |
| | blocked summary | veridical | M = 16.8% [10.3% 23.3%] | M = 30.9% [24.0% 37.9%] | M = 20.5% [15.8% 25.1%] | M = 31.7% [20.4% 43.1%] |
| | | noisy | M = 14.3% [7.1% 21.5%] | M = 40.5% [31.2% 50.0%] | M = 20.7% [15.3% 26.1%] | M = 24.4% [13.0% 35.9%] |
| | rolling summary | veridical | M = 11.1% [6.5% 15.7%] | M = 32.8% [24.5% 41.1%] | M = 21.0% [16.6% 25.5%] | M = 35.0% [22.4% 47.5%] |
| | | noisy | M = 9.8% [4.3% 15.3%] | M = 43.2% [32.0% 54.4%] | M = 21.9% [16.8% 27.0%] | M = 25.1% [13.0% 37.2%] |
