## Supplementary material for "The role of feedback for sensorimotor decisions under risk": Suppl. Fig. S2

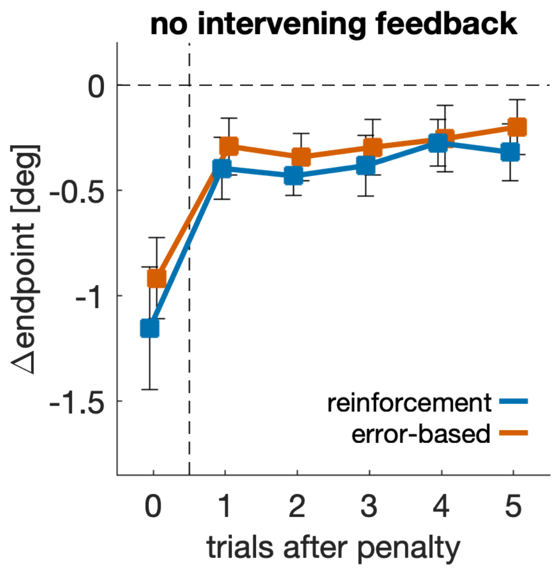


**Suppl. Fig S2. Post-penalty behavior without intervening feedback.** Endpoints relative to the individual mean after encountering a penalty. Data from the blocked summary condition with no feedback in between the penalty trial (T0) and the first five trials preceding the penalty (T1 to T5). The figure/analysis is based on 999 penalties out of the 1163 penalties underlying the blocked summary panel in Figure 5A. Error bars are 95% confidence intervals of between-participant variability.
