## Supplementary material for "The role of feedback for sensorimotor decisions under risk": Suppl. Fig. S1

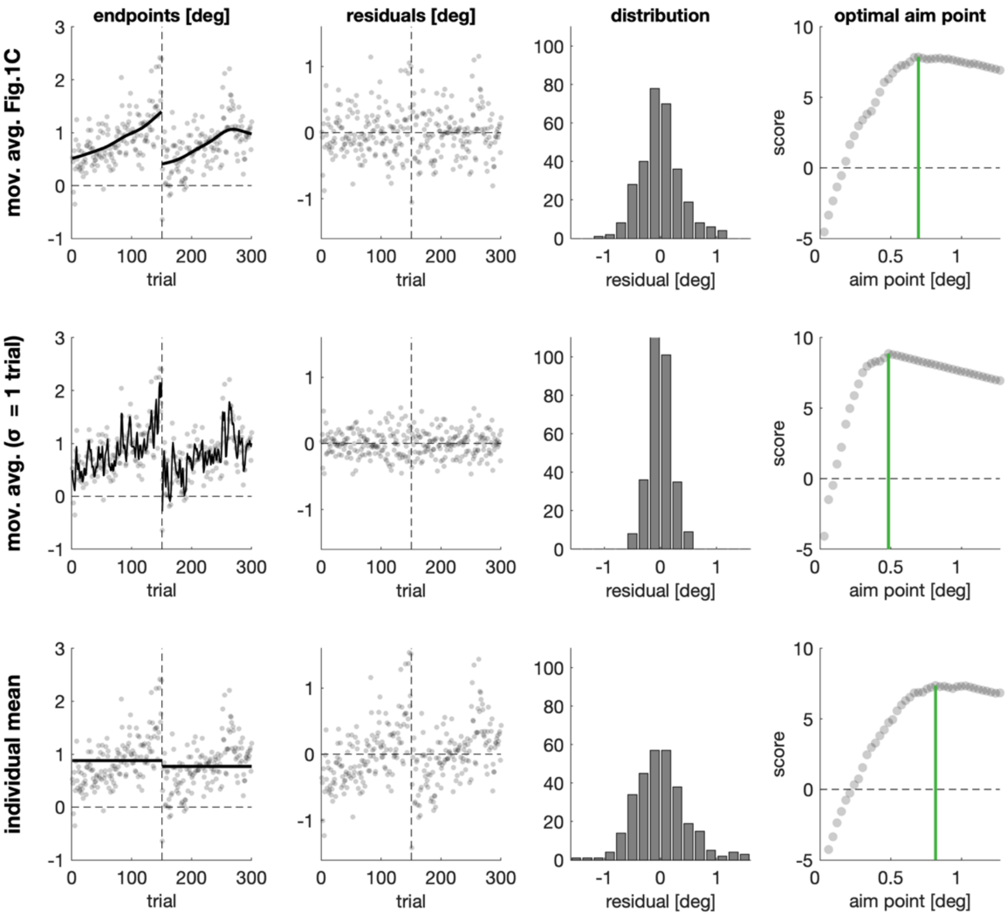


**Suppl. Fig S1. Region of optimality.** Approach for computing the region of optimality based on the individual data shown in Figure 1C. Each row shows the computation of an optimal aim point (green line in right column). Either derived using the moving average depicted in Fig. 1C (top row), the moving average with the smallest sigma parameter (center row) or using the individual mean (bottom row). Whereas the moving average with the smallest sigma provides the lowest possible estimate for the optimal aim point, using the individual mean provides the highest estimate. The region of optimality spans the range between the lowest and the highest estimate (i.e. from the green line in the center row to the green line in the bottom row).
